# Stress Granules and FUS condensates recruit functionally distinct sets of RNAs

**DOI:** 10.64898/2026.05.19.726389

**Authors:** Ryoma Yoneda, Masataka Hirasaki, Yuta Terui, Miki Mori, Tomohiro Kaneko, Masafumi Iharada, Kyota Yasuda

**Affiliations:** Division of Biomedical Sciences, Research Center for Genomic Medicine, Saitama Medical University,1397-1 Yamane, Hidaka-shi, Saitama, Japan 350-1241; Department of Clinical Cancer Genomics, Saitama Medical University International Medical Center, 1397-1 Yamane, Hidaka-shi, Saitama, Japan 350-1241; Global Sales Proposal Department, Sales & Solution Center, Life Business, Yokogawa Electric Corporation, 2-3 Hokuyodai, Kanazawa, Ishikawa, 920-0177 JAPAN; Department of Mathematical and Life Sciences, Graduate School of Integrated Sciences for Life, Hiroshima University, 1-3-1 Kagamiyama, Higashi-Hiroshima, Hiroshima 739-8526, Japan; Research Center for the Mathematics on Chromatin Live Dynamics (RcMcD), Hiroshima University, 1-3-1 Kagamiyama, Higashi-Hiroshima, Hiroshima 739-8526, Japan; International Institute for Sustainability with Knotted Chiral Meta Matter (WPI-SKCM^2^), 1-3-2 Kagamiyama, Higashi-Hiroshima, Hiroshima 739-8531, Japan

**Keywords:** granule-resolved transcriptomics, stress granule, FUS, transcriptome, ALS

## Abstract

Cellular homeostasis relies on the organization of RNA and protein into membrane-less organelles. Stress granules (SGs) are well-known hubs for translational repression during cellular stress. Similar condensates formed by RNA-binding proteins such as FUS— mutations in which cause amyotrophic lateral sclerosis (ALS)—remain poorly characterized relative to canonical SGs. Here, we develop Granule-seq, a microcapillary-based granule-resolved RNA sequencing approach, and demonstrate that SGs and FUS condensates are functionally distinct RNA compartments rather than variants of a unified granule class. Granules were individually aspirated, analyzed in small pooled sets, and validated through overlap with known SG components (794/2,759 genes, 28.8%, p < 0.001). Gene Ontology analysis revealed that G3BP-positive SGs sequester transcripts for stress responses and translational control, while FUS condensates are enriched for transcripts essential for synaptic function and neuronal development. Direct comparison showed that 63% of FUS-enriched and 82% of G3BP-enriched genes were granule-specific, with only 493 genes shared. Exploratory sequence analysis revealed modest contributions from compositional features, with functional identity providing primary selectivity. As a proof-of-concept study with limited biological replication, our results suggest that distinct granule identities are established through functionally specialized transcriptomes, a process that may be disrupted in ALS, and provide a framework for understanding RNA sorting.

**Graphic abstract.**
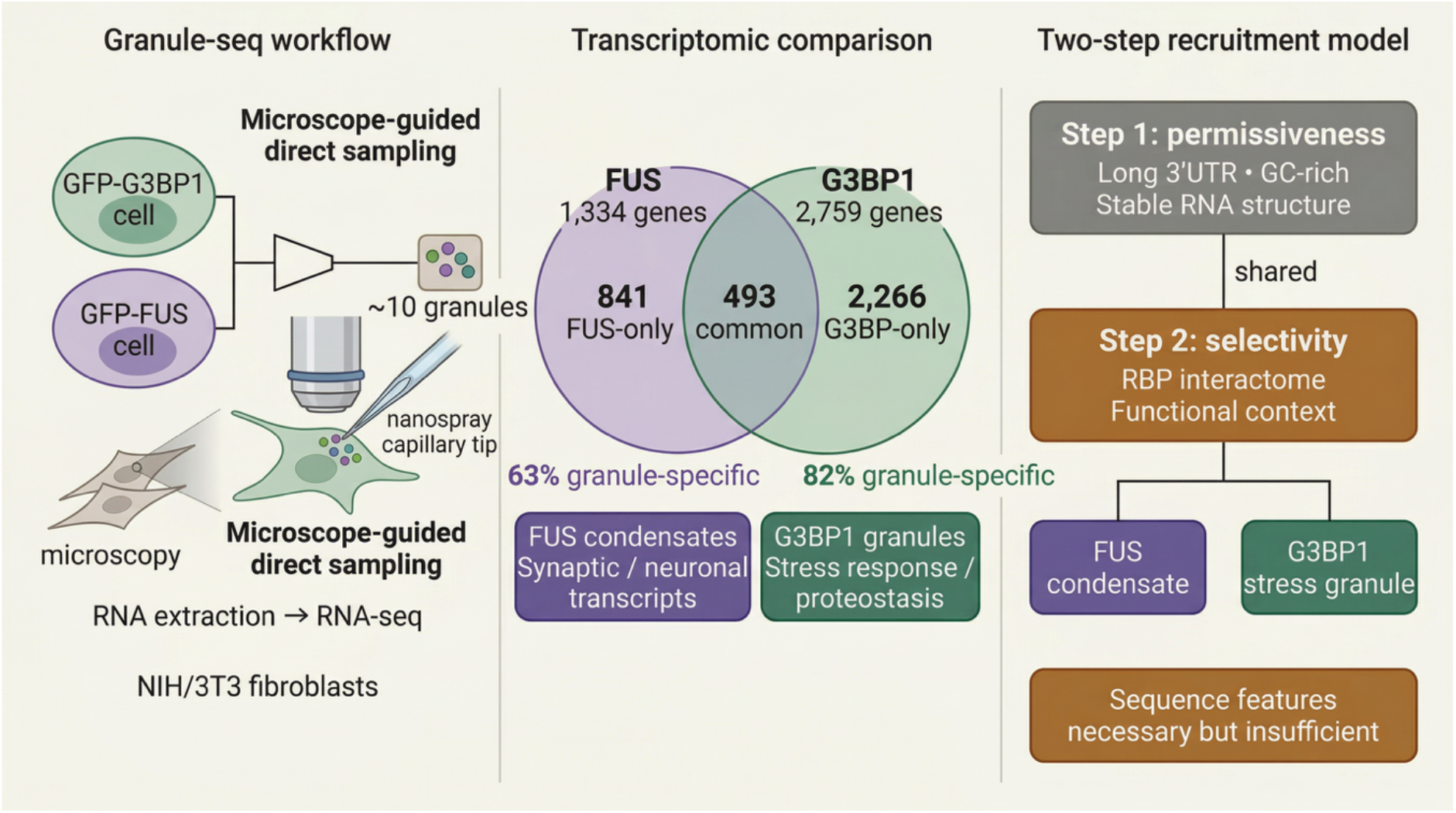
Granule-seq workflow and key findings. (Left) Individual cytoplasmic granules are isolated by microcapillary aspiration, pooled (∼10 granules per replicate), and processed for RNA sequencing alongside total-cell input. (Middle) Venn diagram showing the overlap between FUS condensate-enriched (1,334 genes) and G3BP1 stress granule-enriched (2,759 genes) transcripts. FUS condensates preferentially recruit neuronal and synaptic transcripts; G3BP1 stress granules are enriched for stress-response and proteostasis transcripts. (Right) Two-step recruitment model: shared sequence features (long 3′UTR, GC-rich composition, stable RNA structure) provide permissiveness for granule entry (Step 1), while RBP interactome composition and functional context determine granule-type specificity (Step 2).

## Introduction

The eukaryotic cell is highly organized by a myriad of membrane-less compartments, often referred to as biomolecular condensates, which form through liquid-liquid phase separation (LLPS) to concentrate proteins and nucleic acids^1,2^. Among these, ribonucleoprotein (RNP) granules are prominent hubs for post-transcriptional gene expression, controlling the fate of messenger RNAs (mRNAs) by modulating their translation, storage, and decay^3,4^. RNA itself is not merely a passive client within these granules but often acts as a critical scaffold, with RNA-RNA interactions contributing significantly to granule assembly and integrity^5-7^. Beyond providing multivalent handles for RBPs, specific RNA sequence and structure features can bias partitioning into distinct condensates, producing clientomes with characteristic biological functions. Large-scale annotation frameworks (e.g., KEGG and GO) therefore provide a principled way to read out granule identity from their RNA content^8,9^.

Stress granules (SGs) are canonical RNP granules that assemble in response to a wide range of cellular stresses^10,11^. Their formation is a conserved cellular response triggered by the inhibition of translation initiation, often via phosphorylation of the eukaryotic translation initiation factor 2α (eIF2α), resulting in the accumulation of stalled 48S pre-initiation complexes that serve as seeds for SG nucleation^12,13^. Consequently, SGs are dynamic aggregates of untranslated mRNAs and a host of RNA-binding proteins (RBPs), functioning as sites of mRNA triage and as signaling hubs that orchestrate the stress response^3,14-16^. Accordingly, functional pathway enrichment of SG-associated RNAs has proven informative for distinguishing stress-adaptation programs from other cellular modules^17-19^.

A prominent example of an RBP that forms similar condensates is FUS (Fused in Sarcoma), whose mutations are a direct genetic cause of amyotrophic lateral sclerosis (ALS)^20,21^. LLPS-state FUS engages a protein and RNA interactome that is partially distinct from its soluble state, suggesting that FUS-positive condensates could recruit specialized RNA cohorts in cells^22-25^. Crucially, cytoplasmic FUS is readily recruited to SGs^26,27^, which have been proposed to act as nexuses for the conversion of dynamic FUS condensates into the irreversible, solid-like aggregates characteristic of disease^28,29^. This has fueled a central debate in the field: does neurodegeneration arise from a loss of nuclear functions of FUS, a toxic gain of function from its cytoplasmic aggregates, or a combination of both^24,30,31^? Indeed, FUS function is itself regulated by its phase-separated state; it was recently shown that LLPS-state FUS interacts with a different set of proteins (e.g., chromatin-related factors) than non-LLPS FUS ^22,32^. Recent work by Mariani et al. demonstrated that ALS-associated FUS P525L mutation reshapes stress granule transcriptomes, shifting from GC-rich to AU-rich composition compared to wild-type FUS, suggesting that disease-associated mutations can alter RNA recruitment patterns within stress granules^33^.

A central unresolved question is whether ALS-associated FUS condensates represent pathological variants of canonical stress granules or constitute functionally distinct RNA compartments. Resolving this question is critical for understanding whether neurodegeneration arises from sequestration of specific neuronal transcripts or from dysregulation of general stress responses. While the sequence-level rules that govern selective recruitment of transcripts into condensates are not well-defined, they appear to be context-dependent, with the initiating stressor, cell-type proteomes, and the balance between ARE-binding and G4-binding proteins all potentially influencing client selection^18,34-37^.

To directly address whether FUS condensates and canonical SGs recruit functionally distinct transcriptomes, we established Granule-seq, a microcapillary-based workflow that individually aspirates and pools small sets (∼10 granules) of cytoplasmic granules for RNA sequencing. Comparison of arsenite-induced G3BP1-positive SGs and ALS-mutant FUS condensates revealed striking functional divergence: G3BP granules were enriched for stress-adaptive transcripts (DNA damage response, translation control, protein quality control), whereas FUS condensates preferentially recruited neuronal RNAs (synapse organization, axon guidance, neuronal development). Sequence analysis revealed that both granule types show partially overlapping sequence tendencies, yet exhibit strikingly different functional profiles, pointing to a two-step recruitment model where sequence features provide permissiveness for granule entry while functional context determines granule-type selectivity.

## Materials and Methods

### Plasmids

Green fluorescent protein (GFP)-tagged mouse FUS, carrying a mutation homologous to human P525L, was used as previously described^23,25^. Mouse G3BP1 coding sequence amplified from NIH/3T3 cDNA was cloned into EGFP-C3.

### Cell culture and granule isolation

NIH/3T3 cells (RIKEN Cell Bank) were grown in Dulbecco’s modified Eagle’s medium (DMEM) (high-glucose with L-glutamine and phenol red) (Gibco) supplemented with 10% FBS (Gibco), penicillin, and streptomycin (Gibco). Plasmid transfection (2 μg per 2.5-cm dish) was performed using Lipofectamine 3000 (Invitrogen) or PEI MAX (Polysciences). Cells were cultured for approximately 24 h after transfection before granule isolation. GFP-FUS P525L formed condensates spontaneously during culture, while GFP-G3BP1-marked SGs were assembled by sodium arsenite treatment (500 nM for 1 h). Granules were detected under fluorescence microscopy and isolated using a microscopy system equipped with glass microcapillaries (SS2200, Yokogawa). Approximately 10 granules were pooled and RNA was extracted with TRIzol (Thermo Scientific). Total cellular RNA from cells with the same treatments was extracted as control input.

### RNA-sequencing

Libraries for RNA-seq were prepared using RNAs extracted as above with SMART-Seq Stranded Kit (634442, Takara Bio) according to the manufacturer’s protocol. Briefly, RNAs were fragmented for 5 min, and library preparation was performed with “Ultra-low Input” protocol (10 cycles of PCR1 and 13 cycles of PCR2). Libraries were validated using Agilent 2100 Bioanalyzer with D1000 screen tape (5067-5582, Agilent). Libraries were then sent to Azenta for QC and NGS by Nova-seq (2×150bp).

### RNA-seq Data processing

Data analysis followed previously reported methods^38^. Transcripts per million (TPM) values were calculated using RSEM and log2-transformed as log2(TPM + 1). Genes showing higher abundance were defined as those with log2 fold change > 1. Gene Ontology (GO) and Kyoto Encyclopedia of Genes and Genomes (KEGG) pathway enrichment analyses were performed using the DAVID web-based tool (https://david.ncifcrf.gov/) with default parameters and the mouse genome as the background. The Benjamini–Hochberg (BH) method was applied to control the false discovery rate (FDR), and a BH-adjusted p-value < 0.05 was considered statistically significant. Scatter plots, heatmaps, and dot plots were generated using R (version 4.5.2). For heatmap visualization, TPM values were subjected to row-wise Z-score normalization.

### qRT-PCR

RNAs collected from granules were also used for cDNA preparation. RNAs were reverse transcribed with SuperScript IV VILO (11756050, Thermofisher) according to the manufacturer’s protocol. qPCR was performed as previously described using primers listed in Table S1^39^.

### Sequence feature analysis

Mouse reference genome and annotations were obtained from Ensembl release 102 (GRCm38/mm10). The genome sequence (Mus_musculus.GRCm38.dna.primary_assembly.fa) and corresponding gene annotation file (Mus_musculus.GRCm38.102.gff3) were used for all analyses.

Transcript annotations including mRNA, five_prime_UTR, ORF, and three_prime_UTR were parsed from the GFF3 file. Transcript IDs (ENSMUST) were linked to gene IDs (ENSMUSG) based on the Parent attribute. For each gene, a representative transcript was selected as the isoform with the longest coding sequence (ORF length).

Genomic coordinates corresponding to 5′UTR, ORF, and 3′UTR were extracted for each representative transcript. Sequences were retrieved from the reference genome using pyfaidx, and exon segments were concatenated according to transcript structure to reconstruct full-length regions. For transcripts on the negative strand, reverse-complement sequences were generated. DNA sequences were converted to RNA sequences by replacing thymine (T) with uracil (U).

UTR sequences shorter than 10 nucleotides were excluded from downstream analyses.

RNA secondary structure was predicted for each region independently using RNAfold from the ViennaRNA package (v2.4.17) with default parameters at 37°C. The minimum free energy (MFE, kcal/mol) was used as a measure of thermodynamic stability.

Sequence features including transcript length, AU content, AU-rich element (ARE) density per kilobase, and minimum free energy (MFE) were calculated for each region (5′UTR, ORF, and 3′UTR).

Data distributions were summarized using the median and interquartile range (IQR). Group comparisons were performed using the Wilcoxon rank-sum test. P-values were adjusted for multiple testing using the Benjamini–Hochberg method. Effect sizes were estimated using Cliff’s delta.

### Motif enrichment and density analyses

#### (1) RBP motif definitions

RNA-binding protein (RBP) motifs were defined using regular expression patterns based on previously reported sequence preferences. The analyzed motifs included TIA1/TIAR, HuR/ELAVL, G3BP1, TDP43, FUS, hnRNPC, FUBP1, PABP, AUF1, KSRP, and Pumilio.

For each transcript region (5′UTR, ORF, and 3′UTR), motif presence (binary) and motif occurrence counts were computed from reconstructed RNA sequences.

#### (2) Length-matched bootstrap enrichment analysis

To account for sequence length differences between groups, motif enrichment was primarily evaluated using a length-matched bootstrap framework.

For each comparison and transcript region, control transcripts were stratified into quantile-based bins according to sequence length. Case transcripts were assigned to these bins using the same boundaries. Within each bin, control transcripts were randomly sampled to match the number of case transcripts, generating a length-matched control set.

This procedure was repeated for 1,000 bootstrap iterations. For each iteration, motif enrichment was assessed using Fisher’s exact test, and odds ratios (ORs) were calculated. The median odds ratio across iterations (OR_median) and the 95% bootstrap interval (OR_ci2.5–OR_ci97.5) were reported.

Empirical two-sided p-values were estimated from the bootstrap distribution as:

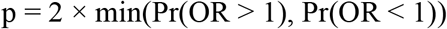

Multiple testing correction across motifs was performed using the Benjamini–Hochberg false discovery rate (FDR) procedure. Motifs with q < 0.05 were considered statistically significant.

#### (3) Length-adjusted logistic regression

As a complementary analysis, motif enrichment was also evaluated using logistic regression. Motif presence was modeled as a binary response variable with case status and log-transformed sequence length as predictors:

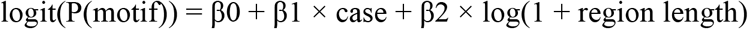

Odds ratios corresponding to the case variable were extracted from the model. In cases where model fitting failed due to complete or quasi-complete separation, Fisher’s exact test was used as a fallback.

P-values were adjusted for multiple testing using the Benjamini–Hochberg FDR procedure.

#### (4) Motif density analysis

Motif density was defined as the number of motif occurrences per kilobase of sequence. Density distributions between case and control groups were compared using a two-sided Mann–Whitney U test.

Median densities and fold changes (case/control) were reported. P-values were adjusted using the Benjamini–Hochberg FDR method, and q < 0.05 was considered statistically significant.

### Gene set definition

Gene sets used for sequence feature analysis and motif enrichment/density analyses were defined as follows. Genes were restricted to those annotated as protein-coding in Ensembl release 113 (Mouse). For condition-specific analyses, genes showing differential abundance between foci and control conditions were identified from RNA-seq data for FUS and G3BP1. From each comparison, the top 200 genes with the largest abundance differences were selected based on ranked differential abundance statistics.

### Reference data sets

The ΔRG-AS reference set comprises 2,920 transcripts defined by Namkoong et al. (2018) as core stress granule-enriched RNAs across multiple stress conditions.

### Statistical analysis

Statistical comparisons were performed using Mann-Whitney U tests for continuous variables (sequence features, motif densities) and Fisher’s exact test for categorical comparisons (gene set overlaps, motif presence/absence). Multiple testing correction was applied using the Benjamini-Hochberg false discovery rate (FDR) method, with q < 0.05 considered statistically significant. Effect sizes for sequence feature comparisons were quantified using Cliff’s delta. All statistical analyses were performed using Python 3.8 with SciPy, NumPy, pandas, and statsmodels libraries. Data visualization was performed using R (version 4.5.2) with ggplot2 and ComplexHeatmap packages.

## Results

### Establishment, validation, and individual transcriptomic profiling

To compare the transcriptomes of stress granules (SGs) and FUS condensates using granule-level sampling, we established Granule-seq, a microcapillary-based workflow in which individually aspirated cytoplasmic granules were collected in small pools for RNA sequencing (Fig. 1A). In NIH/3T3 cells, arsenite-induced G3BP1-positive SGs and spontaneously formed GFP-FUS condensates (carrying the ALS-associated P525L mutation) were visualized by fluorescence microscopy and individually aspirated using glass microcapillaries (Fig. 1B). RNA extracted from pools of approximately 10 granules yielded high-quality material with sharp ribosomal RNA peaks and minimal degradation, suitable for low-input RNA-seq library construction.

**Figure 1.**
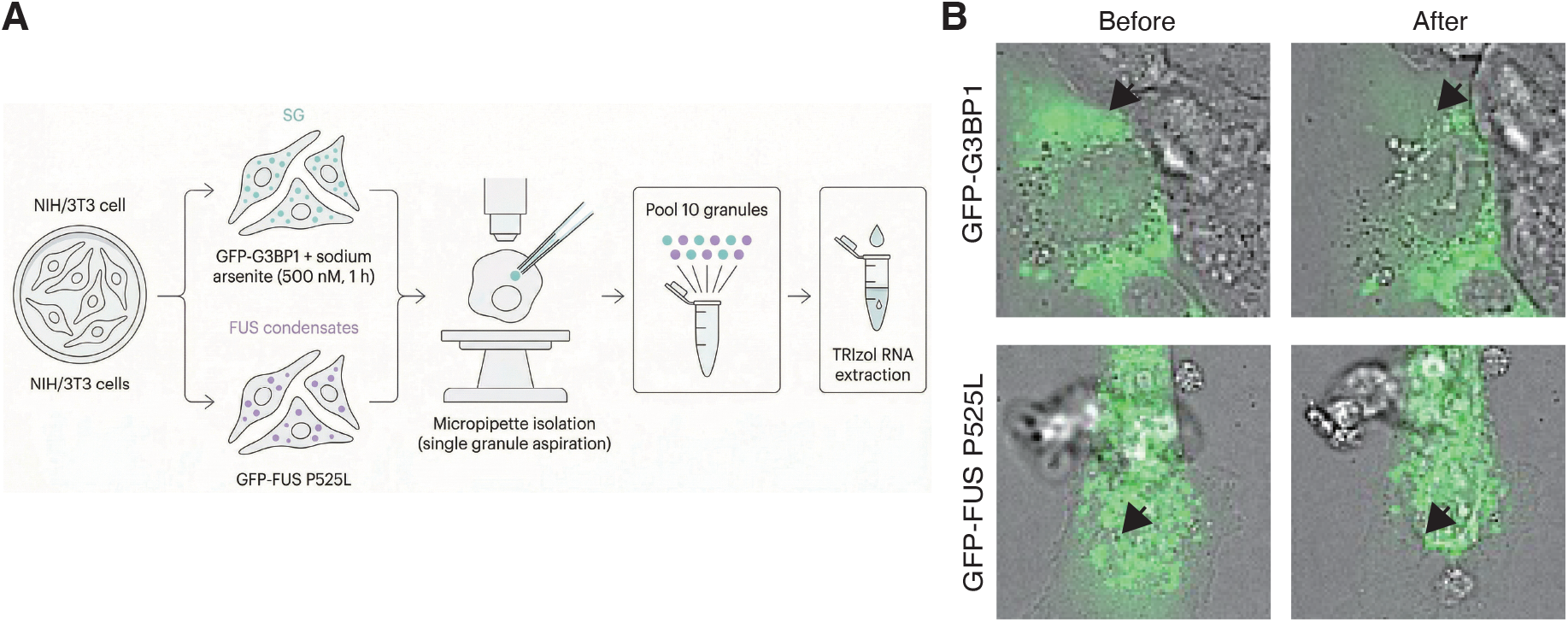
Establishment and validation of the Granule-seq workflow. (A) Schematic of the Granule-seq method. Stress granules (SGs) were induced in arsenite-treated NIH/3T3 cells (G3BP1-positive), and FUS condensates were induced by expression of ALS-mutant GFP-FUS P525L. Individual granules were isolated under fluorescence microscopy using glass microcapillaries, collected in small pooled sets, and processed for RNA-seq alongside total-cellular (input) RNA. (B) Representative images showing granule isolation. Left panels show cells with fluorescent granules before aspiration; right panels show the same fields after individual granule capture. Arrows indicate targeted granules. Scale bars, 10 μm.

To investigate transcriptional changes accompanying FUS and G3BP1 foci formation, we compared gene expression levels between control and foci conditions. Comparative transcript profiling identified 1,334 genes showing higher abundance in FUS condensate samples (log_2_FC ≥ 1) and 2,759 genes in G3BP stress granules relative to whole-cell input (Fig. 2A-B, Table S2a-b). To validate the method, we compared our transcriptomes with a published bulk stress granule dataset^19^. As illustrated in Venn diagrams (Fig. 2C-D), substantial overlap was observed: 374 of 1,334 FUS-enriched genes (28.0%) and 794 of 2,759 G3BP-enriched genes (28.8%) were present in the Namkoong ΔRG-AS reference dataset (Fisher’s exact test, p < 0.001). The substantial overlap with an independent dataset generated using a different methodology validates the technical robustness of our approach.

**Figure 2.**
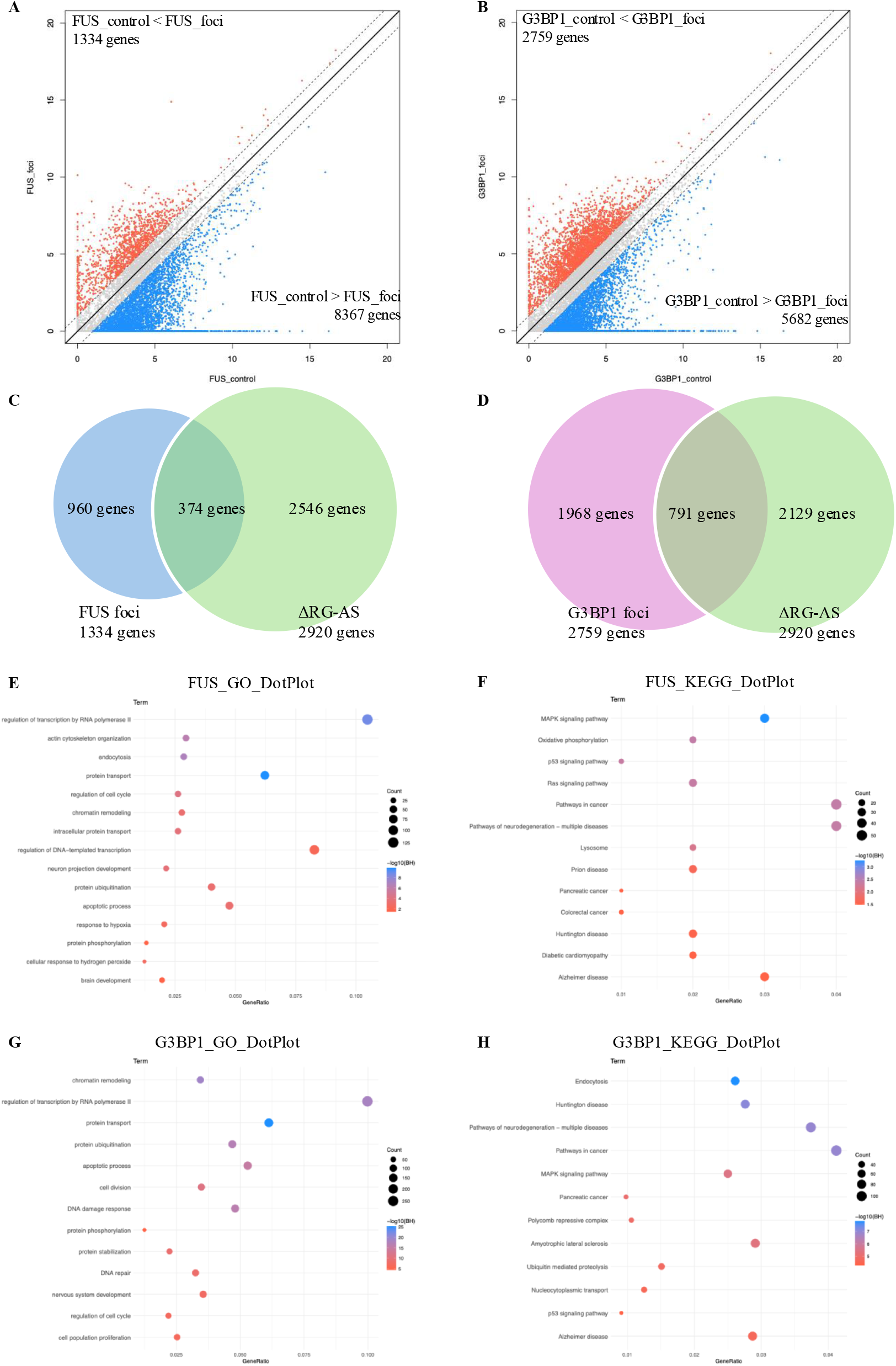
Transcriptomic comparison between control and foci conditions in FUS and G3BP1. (A, B) Scatter plots showing gene expression levels in control and foci conditions for FUS (A) and G3BP1 (B). Genes showing higher abundance in the foci condition (>2-fold) are shown in red, whereas genes showing lower abundance (>2-fold decrease) are shown in blue. The number of genes showing differential abundance is indicated in each panel. (C-D) Venn diagrams showing overlap between Granule-seq datasets and published bulk stress granule transcriptomes (Namkoong et al., 2018, ΔRG-AS). (C) FUS condensates: 374 of 1,334 genes (28.0%) overlap with the Namkoong reference dataset (Fisher’s exact test, p < 0.001). (D) G3BP1 stress granules: 794 of 2,759 genes (28.8%) overlap with the Namkoong reference dataset (Fisher’s exact test, p < 0.001). (E, F) Gene Ontology (GO) biological process (E) and KEGG pathway (F) enrichment analyses of genes showing higher abundance in FUS_foci compared with FUS_control. (G, H) GO biological process (G) and KEGG pathway (H) enrichment analyses of genes showing higher abundance in G3BP1_foci compared with G3BP1_control. In all dot plots, dot size represents gene count and color indicates −log_10_(BH-adjusted p-value).

The larger G3BP set (2.1-fold more genes) is consistent with stress granules serving as broad hubs for stress-responsive transcripts, whereas FUS condensates retained a more selective clientome. Beyond the difference in scale, the two granule types exhibited strikingly divergent functional profiles. Gene Ontology (GO) analysis of FUS-enriched transcripts revealed significant enrichment for biological processes including transcriptional regulation by RNA polymerase II, actin cytoskeleton organization, endocytosis, protein transport, and cell cycle control (Fig. 2E; Table S2c). KEGG pathway analysis identified significant enrichment for MAPK signaling pathway, oxidative phosphorylation, PI3K-Akt signaling pathway, and neurodegenerative disease-related pathways (Fig. 2F; Table S2c). Notably, functional terms related to neuronal development, synapse organization, and axon guidance were selectively enriched among FUS condensate-associated transcripts, consistent with the neuronal bias discussed below (Fig. 2E, F).

In contrast, G3BP1-enriched transcripts showed enrichment for chromatin remodeling, protein transport, ubiquitination, and DNA damage response in GO analysis (Fig. 2G; Table S2d). KEGG pathway analysis revealed significant enrichment for endocytosis, MAPK signaling pathway, ubiquitin-mediated proteasomal degradation, and neurodegenerative disease-related pathways (Fig. 2H; Table S2d).

To further validate the biological relevance of these functional profiles, we compared pathway enrichments between our Granule-seq datasets and the Namkoong bulk stress granule reference. Genes commonly detected in both FUS_foci and Namkoong ΔRG-AS (374 genes) showed concordant enrichment for transcriptional regulation, apoptosis control, and chromatin remodeling (Fig. S2A-B; Table S3a-b). Similarly, genes common to G3BP1_foci and ΔRG-AS (794 genes) exhibited shared enrichment for protein transport, signal transduction, and ubiquitin-dependent degradation pathways (Fig. S2C-D; Table S3c-d). These concordant functional profiles between Granule-seq and bulk methods confirm the biological relevance of our granule-resolved approach. We further validated the method by qRT-PCR on independently prepared samples for selected transcripts, which confirmed enrichment patterns consistent with RNA-seq predictions (Fig. S1).

Together, these results establish Granule-seq as a robust method for granule-resolved transcriptomics and demonstrate that FUS and G3BP1 foci formation influences multiple molecular mechanisms including transcriptional regulation, protein transport, signal transduction, and neurodegenerative disease-related pathways, consistent with distinct functional identities shaped by context-dependent RNA recruitment.

### FUS and G3BP granules recruit largely non-overlapping transcript populations

To directly assess the degree of overlap between FUS and G3BP granule clientomes, we performed comparative analysis of the enriched candidate gene sets. Only 493 genes were common to both granule types, representing 37% of FUS-enriched genes but only 18% of G3BP-enriched genes (Fig. 3A). Conversely, 841 genes (63% of the FUS set) were unique to FUS condensates, and 2,266 genes (82% of the G3BP set) were unique to stress granules. This pronounced asymmetry—with the majority of transcripts in each set being granule-specific—indicates that FUS condensates and stress granules represent functionally distinct RNA compartments rather than minor variants of a unified granule class.

**Figure 3.**
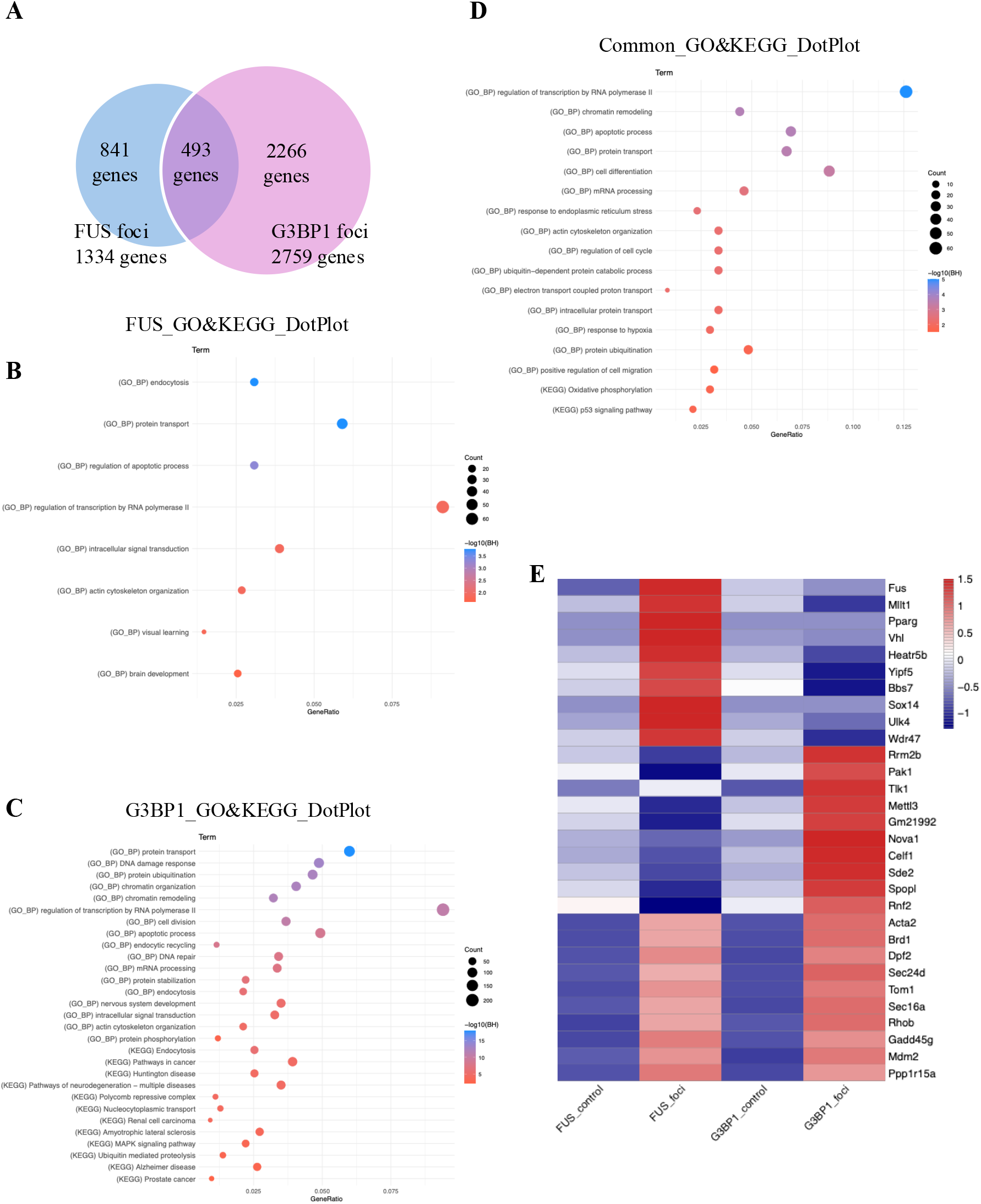
Comparative analysis of FUS- and G3BP1-enriched transcripts. (A) Venn diagram showing the overlap between transcripts showing higher abundance in FUS_foci (1,334 genes) and G3BP1_foci (2,759 genes). A total of 493 genes were commonly enriched, whereas 841 and 2,266 genes were specific to FUS_foci and G3BP1_foci, respectively. (B-D) Gene Ontology (GO) and KEGG pathway enrichment analyses of FUS-specific genes (B), G3BP1-specific genes (C), and commonly enriched genes (D). Dot size represents gene count, and color indicates −log10 (BH-adjusted p-value). (E) Heatmap showing relative expression levels of representative genes across FUS_control, FUS_foci, G3BP1_control, and G3BP1_foci conditions. Transcripts per million (TPM) values were normalized by Z-score transformation as described in the Methods.

The functional profiles of the non-overlapping fractions reinforced this conclusion. FUS-specific transcripts showed significant enrichment for endocytosis, protein transport, transcriptional regulation, and actin cytoskeleton organization (Fig. 3B; Table S2e). In contrast, G3BP1-specific genes exhibited enrichment for DNA damage response, ubiquitination, chromatin remodeling, and cell cycle control (Fig. 3C; Table S2f).

The common set of 493 genes shared between both FUS and G3BP1 foci, though smaller in scale, showed significant enrichment for transcriptional regulation, apoptosis control, protein transport, and signal transduction pathways (Fig. 3D; Table S2g). This suggests that a core layer of universally required transcripts may be partitioned into either granule type depending on cellular context.

Heatmap visualization of representative gene expression patterns showed that common genes exhibited increased expression in both FUS foci and G3BP1_foci conditions, clearly distinguishing them from control conditions (Fig. 3E).

Together, these results indicate that while both granule types share a subset of transcripts, each recruits a functionally distinct clientome, indicating that granule-type identity is determined by context-dependent recruitment rather than shared sequence features alone.

### Sequence features partially distinguish FUS and G3BP1 granule-associated transcripts

To enable balanced comparisons of sequence features, we focused on the top 200 most strongly enriched genes in each condition. To investigate whether the functional divergence between FUS condensates and G3BP1 stress granules could be explained by intrinsic sequence features, we performed a comparative analysis of transcript architecture and composition across the two conditions (Fig. 4; Table S4).

**Figure 4.**
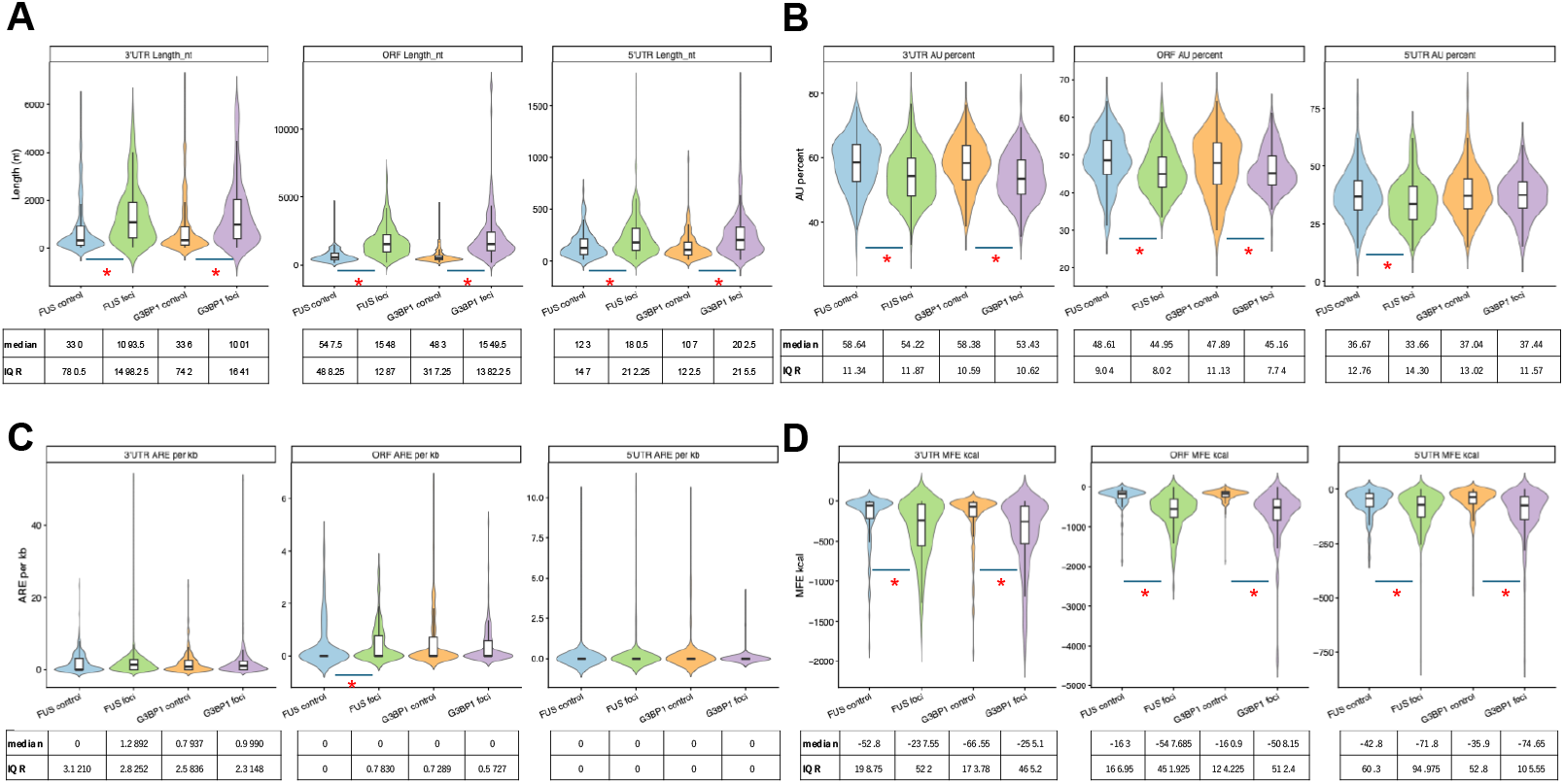
Comparative analysis of mRNA features across FUS and G3BP1 conditions. Violin plots showing the distribution of sequence features in four groups: FUS control, FUS foci, G3BP1 control, and G3BP1 foci. Each violin represents the density of values, with embedded boxplots indicating the median and interquartile range (IQR). (A) Length of 3′UTR, ORF, and 5′UTR (nt). (B) AU content (%) of 3′UTR, ORF, and 5′UTR. (C) ARE (AU-rich element) density per kilobase (ARE per kb) in each region. (D) Minimum free energy (MFE, kcal/mol) of each region. Median and IQR values are shown below each panel. Statistical comparisons were performed using the Wilcoxon rank-sum test with Benjamini– Hochberg correction for multiple testing. Effect sizes were calculated using Cliff’s delta. Asterisks indicate nominal statistical significance (*p < 0.05, Wilcoxon rank-sum test with BH correction); effect sizes were small to modest (Cliff’s delta).

We first examined transcript length across regions. Transcripts enriched in both FUS and G3BP1 foci tended to exhibit longer 3′UTRs compared to their respective control populations, whereas differences in ORF and 5′UTR lengths were more modest (Fig. 4A). These results suggest that extended 3′UTRs may contribute to preferential recruitment into cytoplasmic condensates.

We next analyzed nucleotide composition. AU content showed a modest downward shift in foci-associated transcripts compared to controls across multiple regions (Fig. 4B), indicating that AU richness alone does not fully account for transcript enrichment in either granule type.

We further quantified the density of AU-rich elements (AREs), which are known binding platforms for multiple RBPs. Although ARE density showed region-specific variation, no clear global increase was observed in foci-associated transcripts (Fig. 4C), suggesting that ARE-mediated interactions are not the sole determinant of transcript recruitment.

Finally, we evaluated RNA secondary structure by calculating minimum free energy (MFE). Transcripts enriched in both FUS and G3BP1 condensate showed a trend toward lower MFE values (i.e., more stable predicted structures), particularly within 3’UTRs (Fig. 4D). This trend suggests that RNA structural stability may contribute to condensate partitioning.

Taken together, these results indicate that while sequence features such as 3′UTR length and RNA secondary structure may contribute to granule localization, they only partially explain the distinct transcriptomes of FUS and G3BP1 condensates. These findings support a model in which transcript recruitment is governed by a combination of intrinsic sequence properties and higher-order functional context.

### Motif enrichment analyses reveal distinct RNA-binding signatures in FUS and G3BP1 condensates

To further investigate the molecular determinants underlying selective transcript recruitment, we performed motif enrichment and density analyses for 11 canonical RBP binding motifs across transcript regions (5′UTR, ORF, and 3′UTR; Fig. 5; Table S5).

**Figure 5.**
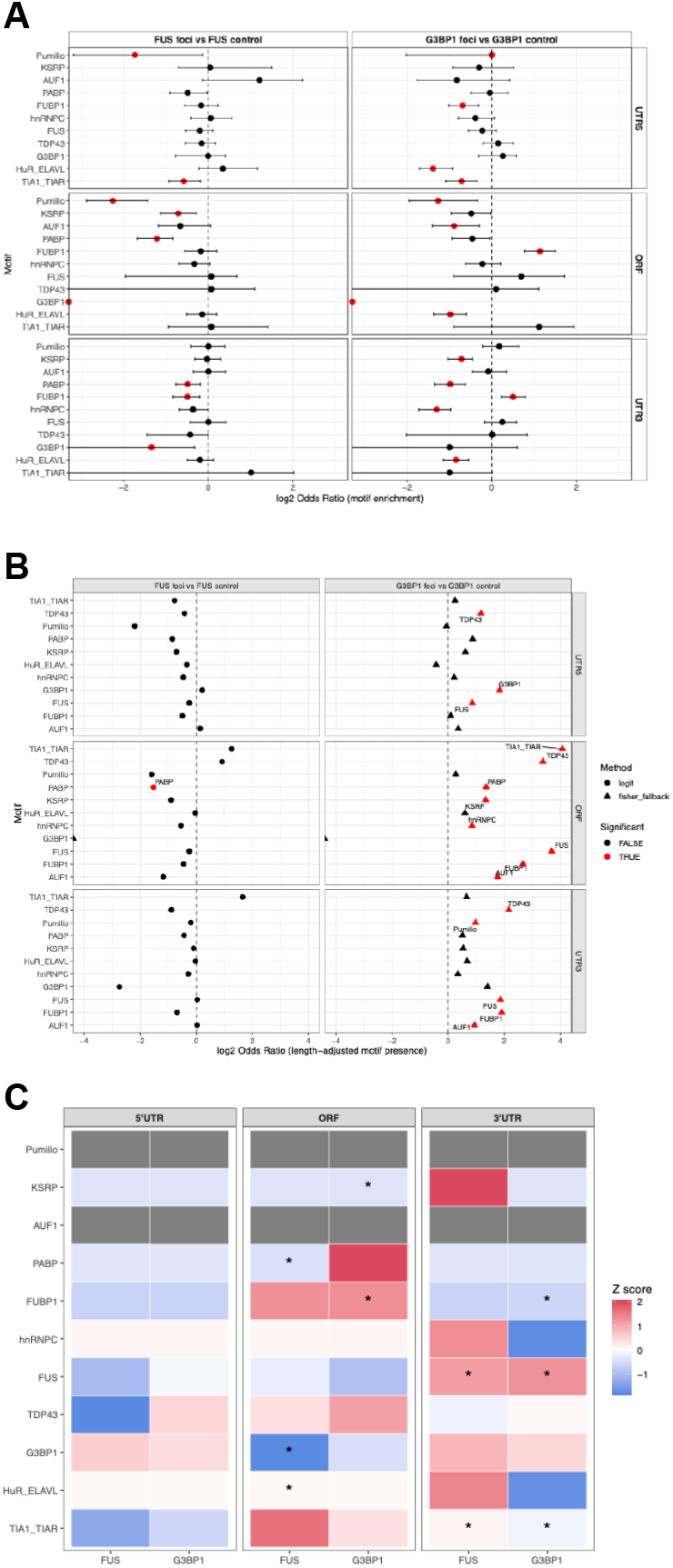
Motif enrichment and density analyses across transcript regions in FUS and G3BP1 conditions. (A) Forest plots showing odds ratios (OR) for motif enrichment in FUS foci versus FUS control (left) and G3BP1 foci versus G3BP1 control (right). Motif enrichment was assessed using logistic regression models with transcript length included as a covariate. Points indicate estimated ORs and horizontal bars represent 95% confidence intervals. Red points indicate statistically significant enrichment after Benjamini–Hochberg correction. (B) Motif density analysis (per kb) comparing foci and control conditions. Each point represents the log_2_ fold change of motif density between groups. Significance was assessed using the Mann–Whitney U test with multiple testing correction. Motifs with significant differences are highlighted in red. (C) Heatmap of standardized motif density (Z-score, row-wise normalization) across 5′UTR, ORF, and 3′UTR regions. Rows represent RNA-binding protein (RBP) motifs and columns correspond to experimental groups. Color intensity indicates relative enrichment (red) or depletion (blue). Black dots indicate statistically significant differences compared to control conditions.

We first assessed motif enrichment using logistic regression with transcript length as a covariate. Several RNA-binding protein (RBP) motifs showed trends toward differential enrichment between foci and control conditions, with distinct patterns observed between FUS and G3BP1 granules (Fig. 5A). Subsets of AU-rich and low-complexity sequence motifs showed increased representation in condensate-associated transcripts, although the direction and magnitude of enrichment varied across regions.

To ensure that these observations were not confounded by transcript length differences, we performed length-matched bootstrap analyses. The results were generally consistent with the regression-based approach, suggesting that the observed motif enrichment patterns were largely consistent across methods (Fig. 5B).

We next examined motif density differences between conditions. Several motifs showed modest changes in density between foci and control transcripts, although the magnitude of these effects was generally modest and varied across motifs and regions (Fig. 5B). These findings indicate that motif abundance contributes to transcript partitioning but does not fully account for the observed differences.

Finally, we integrated motif density across regions and visualized standardized patterns using a Z-score heatmap. This analysis revealed region-specific and granule-type-specific motif signatures (Fig. 5C). Notably, distinct combinations of RBP motifs were enriched in FUS versus G3BP1 condensates, suggesting that each granule type is associated with a characteristic RNA-binding landscape.

Taken together, these results demonstrate that motif-level features contribute to the selective recruitment of transcripts into cytoplasmic condensates. However, similar to global sequence features, motif enrichment alone is insufficient to fully explain the functional divergence between FUS and G3BP1 granules, supporting a multi-factorial model in which combinatorial sequence features and RNA–protein interactions jointly determine RNA partitioning.

## Discussion

In this proof-of-concept study, we directly compared the transcriptomes partitioned into canonical stress granules (SGs) and FUS-positive condensates using a validated granule-resolved RNA sequencing workflow. Three main conclusions emerge.

First, SGs and FUS condensates recruit partly overlapping yet functionally distinct RNA cohorts. Validation through significant enrichment of known stress granule components (794/2,759 genes, 28.8%, Fisher’s exact test p < 0.001) established technical robustness, while functional enrichment analysis revealed that G3BP-positive SGs are biased toward transcripts involved in general stress adaptation, translational control, and protein quality control, whereas FUS condensates are enriched for neurally annotated programs such as synapse organization and axon guidance, consistent with prior observations that condensates can establish distinct client specificities^1,3,17,18,22,40,41^. These functional distinctions were supported by pathway-level analyses: KEGG enrichment mapped FUS-only clients to axon guidance, neurotrophin signaling and spliceosome, whereas G3BP-only clients were biased toward mRNA surveillance, proteasome and autophagy modules (Fig. 2-3; Supplementary Table S2).

Second, this functional segregation was pronounced: 63% of FUS-enriched and 82% of G3BP-enriched genes were granule-specific, with only 493 genes shared between the two sets (Fig. 3A). This substantial divergence indicates that FUS condensates and stress granules represent distinct RNA compartments with specialized biological roles rather than variants of a single entity. The preferential recruitment of neuronal transcripts to FUS condensates, even in non-neuronal NIH/3T3 fibroblasts, is particularly notable and may reflect intrinsic properties of FUS-RNA interactions that become pathologically relevant in neurons where both FUS and its neuronal target transcripts are abundant.

Third, sequence and structural analysis revealed that both granule types tend to recruit transcripts with broadly similar sequence features—relatively long, GC-rich RNAs (Fig. 4A-D)—yet exhibit strikingly different functional profiles (Fig. 2-3). Analysis of canonical RBP recognition motifs revealed an instructive asymmetry: G3BP stress granules showed enrichment for multiple RBP motifs (TDP43, G3BP1, FUS, HuR; Fig. 5C), consistent with their established role as multi-component RBP assemblies^14,35^. In contrast, FUS condensates exhibited limited RBP motif enrichment, including even the canonical FUS binding motif (GGUG), suggesting that LLPS-state FUS employs distinct recruitment mechanisms^22^.

This pattern supports a two-component model of RNA selection: sequence features (length, GC-content, structure) provide permissiveness for granule entry (Step 1), while functional identity and RBP-interactome composition determine granule-type specificity (Step 2). Critically, our comparative analysis reveals that although sequence features such as 3’UTR length and RNA secondary structure appear to contribute to granule recruitment, they are not sufficient to explain the functional divergence between FUS and G3BP1 condensates (Fig. 4-5). This observation—that shared sequence properties support functionally distinct compartments—builds on recent reports showing that simple sequence motifs alone cannot explain granule targeting^17,18,34^, and suggests that functional specialization arises from combinatorial features including RNA structure, protein availability, and cellular context, rather than from sequence determinants alone. This interpretation emphasizes that combinatorial features—including RNA structure, protein availability, and cellular context— collectively determine recruitment efficiency, with granule-type specificity emerging from higher-order functional constraints rather than from intrinsic sequence signatures.

### Integration with recent findings on FUS mutation effects

Our findings complement recent work by Mariani et al. (2024), who demonstrated that ALS-associated FUS P525L mutation reshapes stress granule transcriptomes, shifting from GC-rich to AU-rich composition compared to wild-type FUS. While they compared FUS WT versus mutant within the SG context to understand how disease mutations alter granule properties, we directly compared FUS condensates to canonical G3BP granules to determine whether these represent distinct compartment types. Together, these studies suggest a hierarchical model: FUS mutation alters sequence preference (Mariani et al. 2024), but functional context overrides sequence features to determine granule-type specificity (our study). This hierarchical framework reconciles apparent differences in sequence preferences observed across studies and emphasizes that both mutation state and granule type contribute to RNA clientome specification.

### Reconciling sequence-based observations across studies

Reports emphasizing contributions from G-rich sequences and G-quadruplexes to SG nucleation can be reconciled with our findings by considering context 39,40. The initiating stressor (arsenite engaging eIF2α-dependent arrest in our NIH/3T3 system), the cell-type proteome (relative abundance of G4-binders vs. ARE-BPs), and the assembly stage (nucleation vs. client layering) likely shift client selection. In vitro reconstitution experiments have demonstrated that G-rich RNAs can drive FUS phase separation^42^, and our observation of GC-enrichment in both granule types is consistent with this mechanism. However, the identical GC-enrichment in both FUS and G3BP granules, combined with their divergent functional profiles, is consistent with GC content representing a permissive feature rather than a determinant of granule-type identity. A parsimonious view is that nucleation and client recruitment are partially separable: a more stable core can arise through sequence- or RBP-specific interactions, while a broader shell of client RNAs is subsequently incorporated based on compositional bias and 3’UTR thermodynamics. In our system, pathway-level readouts (GO/KEGG) mirrored these contextual determinants, with G3BP clients aligning to stress-adaptation/proteostasis and FUS clients skewing neuronal.

The observed asymmetry in RBP motif enrichment—multiple motifs in G3BP granules versus limited enrichment in FUS condensates—provides a key mechanistic insight: granule identity is not determined by a universal ‘granule-targeting code’ but rather by context-dependent assembly principles. Stress granules are well-characterized as multi-component RBP assemblies containing TDP43, FUS, HuR, TIA-1, and numerous other RBPs^14,35^, explaining the enrichment of diverse RBP binding motifs. In contrast, FUS P525L condensates form spontaneously without arsenite stress and appear to recruit RNAs through mechanisms less dependent on canonical RBP binding motifs. This suggests that LLPS-state FUS employs recruitment mechanisms distinct from canonical motif-based recognition^22^, a finding that challenges simple sequence-centric models of RNA granule assembly and that FUS binding in vivo is influenced by RNA structure and context beyond simple motif recognition^43^.

### Limitations and future directions

Several considerations qualify the interpretation of these data but do not alter the central conclusions. This proof-of-concept study used single biological replicates for each granule type; future studies with n ≥ 3 will establish statistical robustness for quantitative comparisons and enable assessment of biological variability. Length and base composition covary with folding energetics; joint models that include length, GC/AU ratio and ARE density alongside MFE will better partition direct from mediated effects. MFE estimates reflect equilibrium minima and do not capture co-transcriptional dynamics, RNA modifications, ribosome traffic or in-cell RBP occupancy; orthogonal in-cell probing such as SHAPE- or DMS-guided models would refine the effective energy landscape^44^. FUS condensates were induced by GFP-FUS P525L in fibroblasts, which may under- or over-represent neuronal modules relative to neurons^20^. Finally, enrichment calls and sequence statistics were computed on representative isoforms; incomplete 3’UTR annotations could undercount ARE clusters in a subset of genes.

This framework yields concrete and falsifiable predictions. Reporter 3’UTRs in which AU blocks or ARE clusters are tiled or scrambled should shift granule residency and translational repression in a graded manner, with the largest effects in SG conditions^5,7,10^. Swapping 3’UTRs between FUS-enriched and non-enriched genes should transfer condensate targeting, with motif density and local folding explaining variance^5,7^. Perturbation of ARE-binding proteins is expected to reduce recruitment of ARE-clustered reporters to FUS condensates, whereas modulation of G4-binders should preferentially affect G-rich clients^42,45^. Multivariate models integrating length, AU/GC, ARE density and 3’UTR MFE should quantitatively recapitulate client partitioning, and pathway-level readouts (GO/KEGG) of reporter sets are expected to shift accordingly under ARE-BP or G4-BP perturbations.

Collectively, our results establish Granule-seq as a robust method for granule-resolved transcriptomics and reveal that FUS condensates and stress granules recruit functionally distinct RNA cohorts through mechanisms that extend beyond simple sequence recognition. The observation that shared sequence properties—partially shared sequence features, including transcript architecture and predicted RNA structural tendencies—support functionally divergent compartments supports functional specialization, rather than sequence encoding, as a key determinant of granule identity. The pronounced bias of FUS condensates toward neuronal transcripts, coupled with the known role of FUS mutations in ALS, suggests that disruption of this selective RNA recruitment may contribute to disease pathogenesis. These findings provide a new framework for understanding how RNA granule identities are established through functional specialization and motivate future studies linking 3’UTR-encoded information to phase-separated gene regulation and neurodegenerative disease.

## Data Availability

RNA-seq data have been deposited in DDBJ under accession number PRJDB19728.

## Author Contributions

K.Y. conceived and supervised the project. Y.T., M.M., T.K., and M.I. performed granule isolation experiments. K.Y. performed RNA extraction. R.Y. prepared RNA library and performed qRT-PCR. M.H. and K.Y. performed bioinformatics analysis and statistical testing. K.Y. wrote the first draft of the manuscript. R.Y., M.H., and K.Y. revised the manuscript with input from all authors. All authors read and approved the final manuscript.

## Figure Legends

**Supplementary Figure S1.**
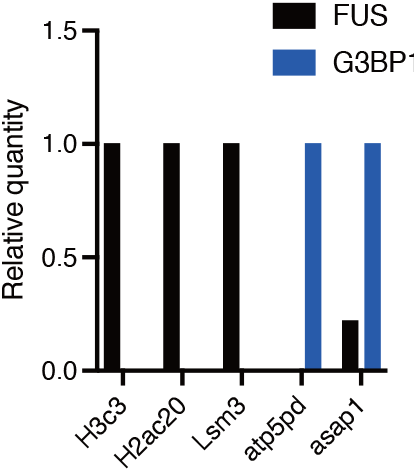
qRT-PCR validation of Granule-seq. qRT-PCR detection of selected transcripts in independently prepared granule samples (n = 3 technical replicates). Bars show relative quantity normalized to input with standard deviation. Transcripts showed enrichment patterns consistent with RNA-seq predictions, validating the accuracy of Granule-seq. Genes: H3c3, H2ac20, Lsm3 (FUS-enriched); atp5pd, asap1 (G3BP-enriched or common).

**Supplementary Figure S2.**
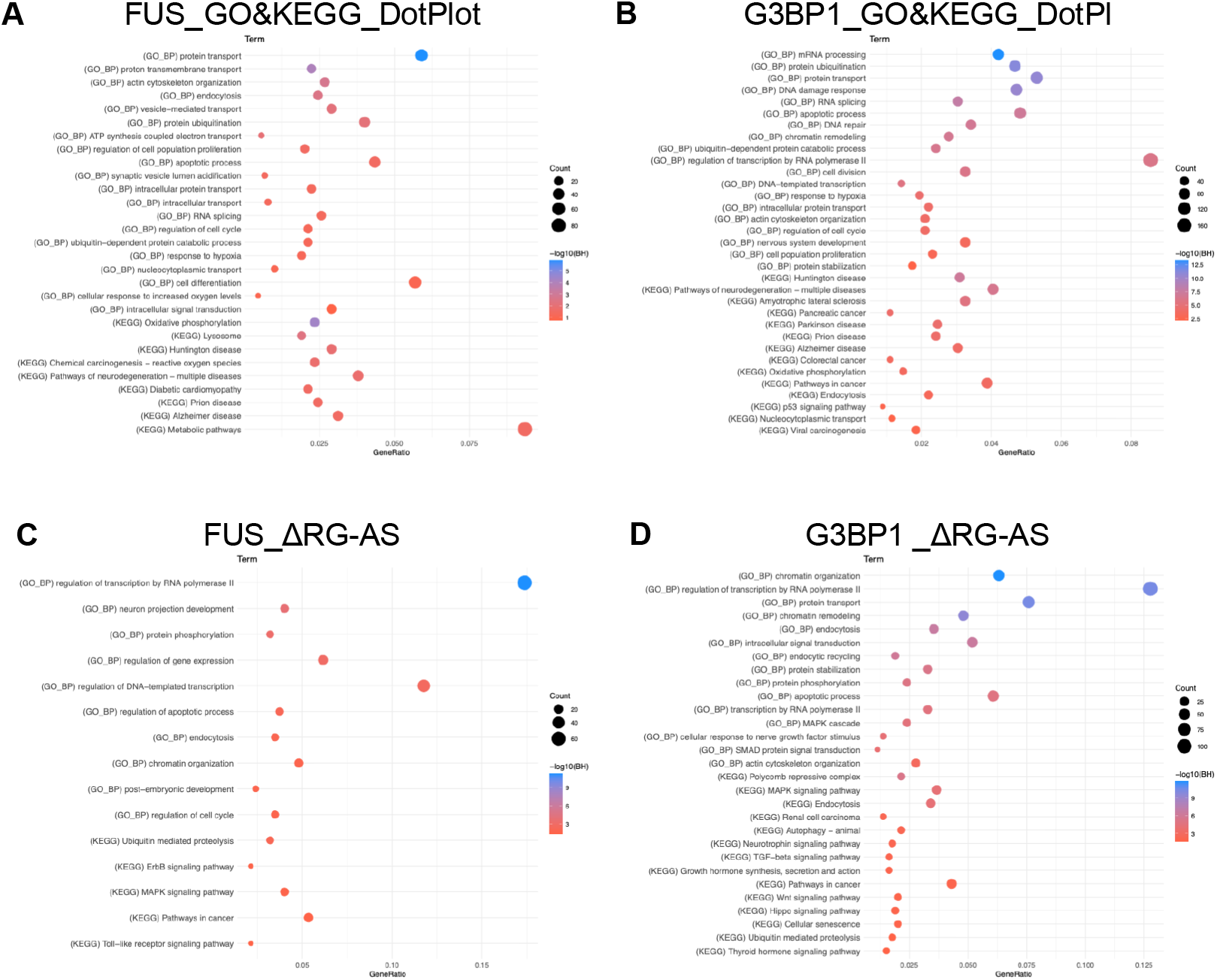
Comparison with Namkoong dataset functional profiles. (A-B) GO and KEGG enrichment comparison between FUS-enriched genes and their overlap with Namkoong ΔRG-AS common transcripts. (C-D) Corresponding analysis for G3BP-enriched genes. Dot size represents gene count, and color indicates −log10 (BH-adjusted p-value).

## Supplementary Tables

**Table S1. Primers used for qRT-PCR validation**.

List of primer sequences used for quantitative RT-PCR validation of selected transcripts identified by Granule-seq. For each gene, the table provides the gene symbol, forward primer sequence, and reverse primer sequence. qRT-PCR was performed on independently prepared biological samples to validate RNA enrichment patterns observed in the RNA-seq analysis (see Figure S1). See Methods for qRT-PCR protocols and analysis procedures.

**Table S2a. Genes showing higher abundance in FUS condensates**.

List of 1,334 genes showing higher abundance in FUS condensate samples relative to FUS_control (log_2_ fold change ≥ 1). For each gene, the table provides Ensembl gene ID, gene symbol, log_2_ fold change, base mean expression, p-value, and FDR-adjusted p-value. Gene type and gene name annotations were retrieved from DAVID; entries marked as N/A indicate genes without available annotation.

**Table S2b. Genes showing higher abundance in G3BP1 stress granules**.

List of 2,759 genes with higher abundance relative to G3BP1_control (log_2_ fold change ≥ 1). For each gene, the table provides Ensembl gene ID, gene symbol, log_2_ fold change, base mean expression, p-value, and FDR-adjusted p-value.

**Table S2c. GO biological process and KEGG pathway enrichment for FUS-enriched genes**.

Functional annotation of the 1,334 FUS-enriched genes using DAVID functional annotation tool. The table lists significantly enriched GO biological processes and KEGG pathways (FDR < 0.05), with term description, gene count, fold enrichment, p-value, and FDR-adjusted p-value.

**Table S2d. GO biological process and KEGG pathway enrichment for G3BP1-enriched genes**.

Functional annotation of the 2,759 G3BP1-enriched genes using DAVID functional annotation tool. The table lists significantly enriched GO biological processes and KEGG pathways (FDR < 0.05), with term description, gene count, fold enrichment, p-value, and FDR-adjusted p-value.

**Table S2e. GO biological process and KEGG pathway enrichment for FUS-specific genes**.

Functional annotation of the 841 genes uniquely enriched in FUS condensates (not enriched in G3BP1 stress granules). The table lists significantly enriched GO biological processes and KEGG pathways (DAVID, FDR < 0.05), with term description, gene count, fold enrichment, p-value, and FDR-adjusted p-value.

**Table S2f. GO biological process and KEGG pathway enrichment for G3BP1-specific genes**.

Functional annotation of the 2,266 genes uniquely enriched in G3BP1 stress granules (not enriched in FUS condensates). The table lists significantly enriched GO biological processes and KEGG pathways (DAVID, FDR < 0.05), with term description, gene count, fold enrichment, p-value, and FDR-adjusted p-value.

**Table S2g. GO biological process and KEGG pathway enrichment for genes common to both FUS and G3BP1 granules**.

Functional annotation of the 493 genes enriched in both FUS condensates and G3BP1 stress granules. The table lists significantly enriched GO biological processes and KEGG pathways (DAVID, FDR < 0.05), with term description, gene count, fold enrichment, p-value, and FDR-adjusted p-value.

**Table S3a. GO biological process enrichment for genes common to FUS_foci and Namkoong ΔRG-AS**.

Functional annotation of the 374 genes commonly detected in FUS_foci and the Namkoong et al. (2018) bulk stress granule reference dataset (ΔRG-AS). The table lists significantly enriched GO biological processes (DAVID, FDR < 0.05), with term description, gene count, fold enrichment, p-value, and FDR-adjusted p-value. These concordant functional profiles validate the biological relevance of the Granule-seq approach.

**Table S3b. KEGG pathway enrichment for genes common to FUS_foci and Namkoong ΔRG-AS**.

KEGG pathway annotation of the 374 genes commonly detected in FUS_foci and the Namkoong et al. (2018) ΔRG-AS dataset. The table lists significantly enriched KEGG pathways (DAVID, FDR < 0.05), with pathway name, gene count, fold enrichment, p-value, and FDR-adjusted p-value.

**Table S3c. GO biological process enrichment for genes common to G3BP1_foci and Namkoong ΔRG-AS**.

Functional annotation of the 794 genes commonly detected in G3BP1_foci and the Namkoong et al. (2018) ΔRG-AS dataset. The table lists significantly enriched GO biological processes (DAVID, FDR < 0.05), with term description, gene count, fold enrichment, p-value, and FDR-adjusted p-value.

**Table S3d. KEGG pathway enrichment for genes common to G3BP1_foci and Namkoong ΔRG-AS**.

KEGG pathway annotation of the 794 genes commonly detected in G3BP1_foci and the Namkoong et al. (2018) ΔRG-AS dataset. The table lists significantly enriched KEGG pathways (DAVID, FDR < 0.05), with pathway name, gene count, fold enrichment, p-value, and FDR-adjusted p-value.

**Table S4a. Pairwise statistical comparisons of sequence features across FUS and G3BP1 conditions**.

For each pairwise comparison of the TOP200 gene sets, the table provides feature name, group labels, sample sizes (n), medians, IQR, Wilcoxon rank-sum test p-value, Cliff’s delta effect size, and BH-adjusted q-value.

**Table S4b. Per-gene sequence feature metrics for the top 200 most highly enriched genes in each condition (FUS_control, FUS_foci, G3BP1_control, G3BP1_foci)**. For each gene, the table provides Ensembl gene ID and length (nt), AU content (%), ARE density (per kb), and MFE (kcal/mol) for 5′UTR, ORF, and 3′UTR regions.

**Table S5a. RBP motif enrichment by length-matched bootstrap analysis (1**,**000 iterations)**. For each motif–region–comparison combination, the table provides OR_median, 95% bootstrap CI, empirical p-value, and BH-adjusted q-value.

**Table S5b. RBP motif enrichment by length-adjusted logistic regression**. For each motif–region–comparison combination, the table provides odds ratio, p-value, analytical method (logit or Fisher fallback), and BH-adjusted q-value.

**Table S5c. RBP motif density analysis across transcript regions**. For each motif– region–comparison combination, the table provides median density (per kb) in case and control groups, fold change, Mann–Whitney U test p-value, and BH-adjusted q-value.

